# Diurnal Modulation of Persistent Inward Current Contribution to Spinal Motor Neuron Behaviour

**DOI:** 10.1101/2025.07.30.667593

**Authors:** Bastien Bontemps, Thomas Cattagni, Simon Avrillon, François Hug

## Abstract

Despite the critical role of persistent inward currents (PICs) in modulating motor neuron output, and thus neuromuscular performance, it remains unknown whether their contribution to motor neuron discharge behaviour varies throughout the day. This study aimed to determine whether PIC-related effects on motor neuron activity during submaximal dorsiflexion tasks differ between the early morning and late afternoon. Eighteen healthy adults (4 females; 27.4±5.6 years) performed triangular isometric contractions at two time-points: early morning (7:00–8:30 a.m.) and late afternoon (5:00–7:30 p.m.). Two conditions were tested: (1) a *relative* condition, where the target force corresponded to 40% of the maximal voluntary force (MVF) measured during that session, and (2) an *absolute* condition, where the target force was 40% of the MVF recorded during the first session. High-density surface electromyography signals were recorded from the *tibialis anterior* and decomposed into motor unit spike trains. The prolongation effect of PICs, estimated *via* ΔF, was significantly greater in the late afternoon in both the relative and absolute force conditions. The amplification effect of PICs, estimated by the acceleration phase of the discharge trajectory, was also higher in the late afternoon, but only in the relative force condition. No time-of-day differences were found for brace height, while attenuation was reduced in the late afternoon in the relative force condition. Collectively, these findings provide evidence for a diurnal modulation of the influence of PICs on motor neuron discharge behaviour, likely mediated by reduced inhibitory input in the late afternoon rather than by changes in neuromodulatory drive.

**Key points:**

‐ Human neuromuscular performance fluctuates throughout the day, with higher force output typically observed in the late afternoon.
‐ While circadian effects on peripheral determinants of muscle function (e.g., muscle– tendon properties, metabolic efficiency) are well documented, the neural mechanisms underlying these diurnal variations remain poorly understood.
‐ Persistent inward currents (PICs) play a critical role in regulating motor neuron behaviour, yet their potential diurnal modulation has not been previously investigated.
‐ Our results provide novel evidence for a diurnal modulation of the influence of PICs on motor neuron discharge behaviour, likely mediated by reduced inhibitory input in the late afternoon rather than by changes in neuromodulatory drive.
‐ These findings suggest that intrinsic motor neuron excitability (and therefore spinal gain modulation) fluctuates across the day.

## INTRODUCTION

Human neuromuscular performance fluctuates throughout the day. For example, maximal force production typically peaks in the late afternoon compared to the morning (Douglas et al., 2021; Knaier et al., 2022). Circadian influences on peripheral determinants of muscle force, such as muscle-tendon mechanical properties and metabolic efficiency, are well documented (Martin et al., 1999; Pearson & Onambele, 2006; Winget et al., 1985). However, the contribution of neural mechanisms to these diurnal fluctuations in force production remains poorly understood (Hirono et al., 2024; Lagerquist et al., 2005; Tamm et al., 2009).

Alpha motor neurons constitute the final common pathway for movement control (Sherrington, 1911). These neurons integrate ionotropic excitatory and inhibitory inputs *via* nonlinear processes that are shaped by intrinsic membrane properties, notably persistent inward currents (PIC) (Heckman et al., 2004). These voltage-dependent depolarizing currents, mainly mediated by dendritic L-type calcium and persistent sodium channels, amplify synaptic inputs (PIC amplification) and sustain motor neuron firing even when excitatory ionotropic inputs decline (PIC prolongation) (Binder et al., 2020; Heckman et al., 2004). As such, PICs play a crucial role in regulating motor output (Heckman & Enoka, 2012)

The activation of PICs is strongly modulated by neuromodulatory inputs, particularly serotoninergic and noradrenergic projections originating from the brainstem (Elliott & Wallis, 1992). While studies on brain tissue samples have demonstrated a pronounced circadian rhythmicity in neuromodulator concentrations, with serotonin levels peaking from late afternoon to nighttime (Carlsson et al., 2007; Mateos et al., 2008; Matheson et al., 2015), it remains unclear whether a similar pattern exists at the level of the spinal cord. Notably, a recent study in older adults reported no evidence of diurnal variation in spinal serotonin levels (Anand et al., 2024), leaving it unclear whether neuromodulatory inputs that influence spinal motor neurons PICs fluctuate throughout the day.

Importantly, PICs are also sensitive to inhibitory inputs (Bui et al., 2008; Hyngstrom et al., 2008; Kuo et al., 2003), which can significantly reduce or suppress their activation. Human studies have suggested that cortical inhibition is reduced in the evening compared to the morning (Lang et al., 2011; Sale et al., 2008), although conflicting findings have been reported (Brangaccio et al., 2024). In addition, previous findings support the notion of reduced spinal inhibition in the evening (Lagerquist et al., 2005). Collectively, these time-of-day differences in inhibitory mechanisms, at both cortical and spinal levels, may also contribute to diurnal variations in PIC activation.

Despite the critical role of PICs in modulating motor output, it remains unknown whether their activation and effects on motor neuron behaviour varies throughout the day, which may in turn contribute to diurnal variations in neuromuscular performance. The present study aimed to determine whether PIC-related contribution to motor neuron activity differ between the early morning and late afternoon, two time points corresponding to the bathyphase and acrophase of neuromuscular performance, respectively (Davison, 2022). Using decomposition of electromyographic (EMG) signals, we identified the discharge patterns of a large sample of motor units in the *tibialis anterior*. We used complementary methodological approaches to estimate the respective contributions of neuromodulatory and inhibitory inputs to motor unit discharge behaviour. We hypothesized that estimates of PIC contribution to motor unit activity would be greater in the late afternoon, primarily reflecting lower inhibitory inputs to spinal motor neurons.

## METHODS

### Participants and ethical approval

Twenty-two healthy adults volunteered for the study. Of these, eighteen participants (4 females; age 27.4 ± 5.6 years; height 174.8 ± 6.6 cm; body mass 70.1 ± 11.4 kg) yielded successful motor-unit decomposition at all time points and were therefore included in the final analyses. Based on the Morningness-Eveningness Questionnaire (Horne & Ostberg, 1976), participants’ chronotypes were categorized as “moderate morning” (n=3; 2 females; score: 53.4 ± 9.5), “intermediate” (n=12, 2 females; score: 49.3 ± 5.3), or “moderate evening” type (n=3; score: 39.3 ± 1.6), with an overall score of 49.5 ± 8.8. None of the participants reported lower limb injuries in the six months preceding testing. Participants were instructed to refrain from caffeine consumption and moderate-to-intense physical activity for 24 hours prior to the evaluations. In addition, they were asked to avoid any physical activity before arriving at the laboratory for the “morning” session, to minimize its potential impact on serotonin concentration (Chaouloff, 1997). The procedures were approved by the ethics committee (CPP Sud-Est V; 2023-A02544-41), and were performed according to the Declaration of Helsinki, except for registration in a database. All participants were fully informed of any risks or potential discomfort associated with the procedures before providing written informed consent.

### Experimental design and set up

Participants attended the laboratory on two separate occasions: once in the morning (between 7:00 and 8:30 a.m.) and once in the late afternoon (evening) on a different day (between 5 and 7:30 p.m.). The two sessions were scheduled 2 to 3 days apart. For the morning session, the average wake-up time was 5:58 ± 0:12 a.m., with a mean interval of 76 ±□12 minutes between waking and the start of data collection. The average onset of data collection was 8:13□±□0:19 a.m. for the morning session, and 6:07□±□0:32 p.m. for the late afternoon session. Room temperature was kept constant across visits (20.0 ± 1.0 °C).

Each visit followed identical experimental procedures. Participants were seated comfortably in a custom-built chair connected to an ankle ergometer (DinamometroGC, OT Bioelettronica, Torino, Italia). The right limb was securely attached to the footplate. Participant’s position was adjusted to maintain an ankle angle of 0° (neutral position), a hip angle of 80° (with full extension defined at 180°), and a minimum knee flexion of approximately 10° (with full extension at 0°). Dorsiflexion force was sampled at 2048 Hz (EMG-Quattrocento, 400-channel EMG amplifier; OT Bioelettronica, Torino, Italy).

### Experimental protocol

#### Maximal voluntary contraction force

Participants first completed a standardized warm-up consisting of 5-second dorsiflexion contractions at progressively increasing intensities, ranging from 30% to 90% of perceived maximal effort, in 10-20% increments. Contractions were separated by a 20-second rest period. After a 3-minute recovery, participants performed two maximal voluntary dorsiflexion efforts, each lasting 4 to 5 seconds, with a 2-minute rest between attempts. If the difference between the two trials exceeded 5%, a third trial was performed. The highest value was retained and used as the reference for setting submaximal dorsiflexion targets. During maximal efforts, participants received strong verbal encouragement and real-time visual feedback of the force signal. Peak maximal voluntary force was defined as the maximal value obtained within a 500-ms moving average window.

#### Submaximal isometric contractions

Participants performed a series of submaximal isometric dorsiflexion tasks following a triangular force profile, consisting of a 10-second ramp-up phase followed by a 10-second ramp-down phase. These tasks were performed under two conditions: (1) a *relative* condition, where the target force corresponded to 40% of the maximal voluntary force measured during the same session, and (2) an *absolute* condition, where the target force corresponded to 40% of the maximal voluntary force recorded during the first visit, regardless of whether it was conducted in the morning or late afternoon. These two conditions were tested based on the expectation of higher force output in the late afternoon compared to the morning, allowing us to better isolate the time-of-day differences in PIC activation from those related to force output. Two contractions were performed for each condition (i.e. *relative* and *absolute* force), with a minimum of 90 seconds between contractions. If the produced force deviated noticeably from the expected triangular shape, an additional triangular contraction was performed. To limit this risk, participants were first familiarized with the triangular contractions.

#### High-density surface electromyographic recordings

High-density surface electromyograms (HDsEMG) were recorded from the *tibialis anterior* (TA) of the right leg using four adhesive grids, each consisting of 64 electrodes (256 electrodes in total), with a 4-mm inter-electrode distance (13 × 5 gold-coated electrodes, with one electrode absent in a corner; GR04MM1305, OT Bioelettronica, Torino, Italy). After identifying the muscle borders with ultrasound imaging, the skin was shaved and cleaned with an abrasive paste (Nuprep, Weaver and Company, USA). Three grids were placed in series along the tibia, while the fourth was positioned adjacent to them over the widest part of the muscle belly. This configuration was chosen to maximize the motor unit yield (Caillet et al., 2023). Grids were held in place on the skin using double-sided adhesive foam layers, and the skin-electrode contact was made by filling the cavities of the adhesive layers with conductive paste (SpesMedica, Battipaglia, Italy). The placement of the grids was carefully marked on the skin using a black marker to facilitate accurate repositioning during the second visit. A reference electrode (5×5 cm; Kendall Medi-Trace; Covidien, Ireland) was placed over the right tibia (distal area), and a strap electrode dampened with water (ground electrode) was placed around the left ankle. The HDsEMG signals were recorded in a monopolar mode, bandpass filtered (10–500 Hz), and digitized at a sampling rate of 2048 Hz using a multichannel acquisition system (EMG-Quattrocento; 400-channel EMG amplifier; OT Bioelettronica, Italy). The data were acquired using OT BioLab+ software (OT Bioelettronica, Torino, Italy).

#### Decomposition of electromyographic signals

EMG signals were decomposed using a source separation algorithm (Negro et al., 2016) implemented in the MUedit software (Avrillon et al., 2023). This process enabled the extraction of individual motor unit pulse trains, from which discharge times were identified. All identified motor unit firings were visually inspected and manually edited when necessary. This procedure was performed according to the guidelines described by Del Vecchio et al. (2020) and was demonstrated to be highly reliable across operators (Hug et al., 2021). Decomposition was performed independently on each electrode grid, and duplicate motor units were identified. Specifically, discharge times occurring within a 0.5 ms interval were considered common between motor units, and those sharing more than 30% of their discharge times were classified as duplicates (Holobar et al., 2010). In such cases, the motor unit spike train with the lowest coefficient of variation was retained.

### Motor unit discharge characteristics

Each motor unit instantaneous discharge rate was smoothed using support vector regression to obtain a continuous estimate of discharge rate, as described in Beauchamp et al. (2022). Simultaneously, the force signal was interpolated to align with the time produced by the support vector regression and then smoothed using a moving average filter with a window width of 1 second (i.e., equivalent to the sampling frequency), ensuring accurate temporal alignment with the discharge rate data. The peak discharge rate was defined as the highest value obtained from the smoothed discharge rate profile. Additionally, the discharge rate at recruitment was defined as the highest discharge rate within the first six interspike intervals following motor unit recruitment (Martinez□Valdes et al., 2020). The recruitment threshold was expressed as the relative force at which the motor unit began discharging, defined as the occurrence of at least three spikes within a 0.7-second window.

### Paired motor unit analysis

The PIC prolongation effect was estimated using the paired motor unit analysis (Gorassini et al., 2002). Briefly, the smoothed discharge rate of individual motor units was used to calculate the difference in discharge rate of a control unit (a lower-threshold motor unit) between the recruitment and de-recruitment times of a test unit (a higher-threshold motor unit). This difference, referred to as ΔF, is considered proportional to the prolongation effect of PIC (Gorassini et al., 2002; Kiehn & Eken, 1997). ΔF estimates were considered only for motor unit pairs that met the following criteria: (i) the test motor unit discharged for at least 2 seconds, (ii) the test motor unit was recruited at least 1 second after the control motor unit, (iii) the test motor unit was de-recruited at least 1.5 seconds before the control unit to avoid overestimation of ΔF, (iv) the control unit modulated its discharge rate by ≥ 0.5 pps while the test unit was active (Stephenson & Maluf, 2011), and (v) the smoothed discharge rate profile of the two motor units were highly correlated (r ^2^ ≥ 0.7), suggesting a large proportion of common synaptic input (Hassan et al., 2020). ΔF values were derived from 2,317 test motor units, each paired with an average of 22.1 ± 12.5 control units. For each test unit, a single representative ΔF value was calculated by averaging across all pairings, ensuring a robust estimate of PICs.

### Quasi-geometric analyses of motor unit discharge behaviour

We used the quasi-geometric method proposed by Beauchamp et al. (2023) to quantify deviations from linearity in the discharge–torque relationship during the ramp-up phase of the triangular contractions (Figure 1). This approach allows for the assessment of PIC-related synaptic amplification *via* the *acceleration* phase, which reflects the initial increase in discharge rate following recruitment and is attributed to early activation of PIC (Li et al., 2004; Revill & Fuglevand, 2016). To further investigate the underlying mechanisms contributing to the nonlinearity in the discharge–torque relationship — and in particular the relative contributions of neuromodulatory and inhibitory drives — we extracted complementary geometric metrics: *attenuation* and *brace height* (Beauchamp et al., 2023). *Attenuation* corresponds to a reduced rate of increase in discharge rate following the acceleration phase and is thought to reflect the continued rise in PIC conductance, along with the activation of potassium currents (Li & Bennett, 2007). In computational models, a shallower attenuation slope reflects a stronger inhibitory influence on motor neurons (Beauchamp et al., 2023; Chardon et al., 2024). *Brace height* quantifies the peak deviation of the discharge rate trajectory from a linear fit between recruitment and peak discharge. This metric has been shown in computational models to be highly sensitive to neuromodulatory input: a higher brace height reflects stronger monoaminergic drive and enhanced PIC conductance, leading to a more pronounced convexity in the discharge rate trajectory (Beauchamp et al., 2023; Chardon et al., 2024).

**Figure 1.**
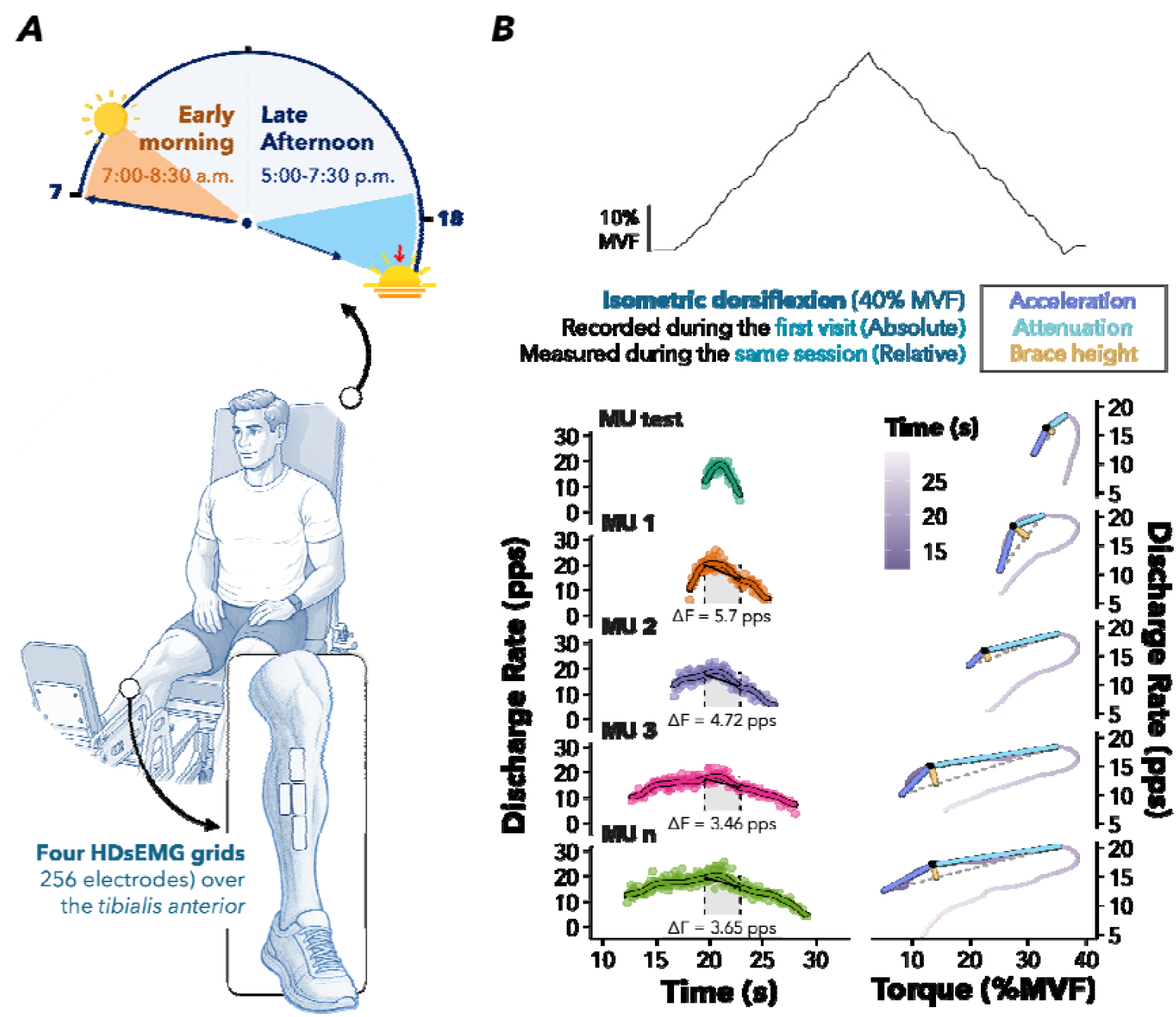
Experimental setup and analyses. (**A**) Overview of the experimental protocol, including time-of-day sessions (early morning: 7:00–8:30 a.m.; late afternoon: 5:00–7:30 p.m., top panel) and participant setup (bottom panel). Participants were seated in a custom-built chair with the ankle attached to a dynamometric ergometer. Four high-density surface EMG (HDsEMG) grids (256 electrodes in total) were positioned over the *tibialis anterior* muscle. (**B**) Example from one participant (subject #16) of the force profile during a triangular contraction at 40% of maximal voluntary force (MVF, top panel). The paired motor unit analysis (bottom left) was used to estimate the PIC-related prolongation effect. ΔF estimates for few motor units are illustrated as the difference in discharge rate of a control unit (lower-threshold MU, shown in in orange, purple, pink, and green) at the recruitment and de-recruitment times of a test unit (higher-threshold MUs, shown in cyan). The geometric approach (bottom right) analysed the discharge rate–torque relationship of these motor units to extract three metrics: acceleration, attenuation, and brace height.

To compute these three metrics, we first fitted a straight line to the smoothed discharge rate trajectory of each motor unit from the time of recruitment to peak discharge rate (Figure 1). The maximum perpendicular distance between this linear fit and the actual smoothed discharge rate was defined as *brace height*. To account for differences in recruitment thresholds across motor units and their associated differences in discharge rate, brace height was normalized to the height of a right triangle whose hypotenuse corresponds to the linear segment from recruitment to peak discharge rate (Beauchamp et al., 2023). Motor units exhibiting a negative brace height prior to normalization were excluded from further analysis. Additionally, the discharge rate trajectory was segmented into two temporal phases: an *acceleration* phase, extending from the first discharge to the point at which brace height occurred, and an *attenuation* phase, extending from the brace height point to the peak discharge rate (see above). Motor units were excluded if they exhibited a negative slope during the acceleration phase or if peak discharge occurred after the peak of the torque profile. Furthermore, because the quasi-geometric method relies on segmenting the discharge rate trajectory into acceleration and attenuation phases, a brief attenuation period may reflect an incomplete trajectory (i.e. when the true peak discharge rate has not yet been reached), potentially introducing artifacts in the estimation of brace height. To mitigate this, motor units were excluded if the attenuation phase lasted less than an empirically determined 2-second threshold. This criterion effectively corrected for an over-representation of short-duration attenuation phases.

### Statistical analyses

To assess changes in motor unit discharge characteristics across time of day, we first identified and removed outliers at the motor unit level using the 1.5x interquartile range method, applied separately to each metric (i.e., recruitment threshold, peak discharge rate, discharge rate at recruitment, ΔF estimates, brace height, acceleration, and attenuation). For each variable, linear mixed-effects models were fitted with time-of-day (early morning *vs*. late afternoon) as a fixed effect, and either a random intercept or a random slope by participant (ID), depending on model fit [e.g., ΔF ∼ time-of-day + (1 | ID) or ΔF ∼ time-of-day + (time-of-day | ID)]. The random slope model was retained when it significantly improved model fit, as determined by likelihood ratio tests and Akaike/Bayesian Information Criteria (AIC/BIC) comparisons. Separate models were fitted for the *relative* condition (40% of the maximal force achieved during the same session) and the *absolute* condition (40% of the maximal force achieved during the first session), with data from the first session (according to the randomized order) being duplicated and reclassified as ‘absolute’ (rather than analyzing force conditions as an additional fixed effect within the models). Estimated marginal means (EMM ± SE) and pairwise comparisons were computed using the emmeans package, with Satterthwaite-adjusted degrees of freedom. Confidence intervals (95%) were also reported. The normality and homoscedasticity of model residuals were assessed graphically. To assess associations between time-of-day–dependent changes in variables, pairwise correlations were computed between percentage changes from morning to late afternoon. Analyses were performed separately for each force condition. Pearson correlation coefficients were calculated and associated p-values are reported. Non-parametric bootstrap estimates (10,000 resamples) were also computed to assess the robustness of the results. All statistical analyses were performed using R (v.4.4.0; R Core Team 2021, R Foundation for Statistical Computing, Vienna, Austria). The *alpha* level for all statistical tests was set at 5%.

## RESULTS

### Maximal voluntary force and motor unit discharge characteristics

A significant time-of-day effect on maximal voluntary force was observed (p = 0.01), with dorsiflexion force 5.3 ± 6.5% greater in the late afternoon (270 ± 72 N) than in the early morning (257 ± 69 N).

On average, 35.1 ± 18.9 motor units were identified per participant and condition (range: 9– 92). When considering the recruitment thresholds, there was no significant main effect of time-of-day in either the relative force (F(1, 19.74) = 0.098, p = 0.76) or the absolute force condition (F(1, 47.36) = 0.162, p = 0.69). Recruitment threshold values of the identified motor units were in average 17.4 ± 0.4% (95% CI [16.5, 18.3]) and 17.4 ± 0.5% (95% CI [16.4, 18.4]) in the relative and absolute force conditions, respectively. It makes us confident that motor units with comparable intrinsic properties were compared between the two time points.

A significant time-of-day effect was found for peak discharge rate in the relative force condition (F(1, 18.56) = 9.226, p = 0.007), with values significantly higher in the late afternoon (20.8 ± 0.5 pps; 95% CI [19.7, 21.8]) compared to the early morning (19.8 ± 0.5 pps, 95% CI [18.7, 21.0]), corresponding to a relative difference of +4.7% ± 3.7%. In contrast, no significant time-of-day effect was observed for peak discharge rate in the absolute force condition (F(1, 16.50) = 2.074, p = 0.17), with an average peak discharge rate across time points of 20.0 ± 0.5 (95% CI [18.9, 21.1]).

No significant time-of-day effect was observed for discharge rate at recruitment in either the relative (F(1, 15.2) = 1.094, p = 0.31) or the absolute force condition (F(1, 17.96) = 1.407, p = 0.25), with an average peak discharge rate across time points of 10.8 ± 0.2 pps (95% CI [10.4, 11.3]).

### Estimates of persistent inward currents using paired motor unit analysis

We first estimated the prolongation effect of PICs using the paired motor unit analysis technique, which quantifies PIC-mediated discharge hysteresis through the ΔF metric (Gorassini et al., 2002). A significant time-of-day effect was found in the relative force condition (F(1, 17.88) = 6.718, p = 0.019), with ΔF values significantly 0.41 ± 0.21 pps greater in the late afternoon (4.9 ± 0.2 pps; 95% CI [4.5, 5.2]) than in the early morning (4.5 ± 0.1 pps; 95% CI [4.2, 4.7]). This corresponded to a relative increase of +9.3% ± 4.8% (Figure 2).

**Figure 2.**
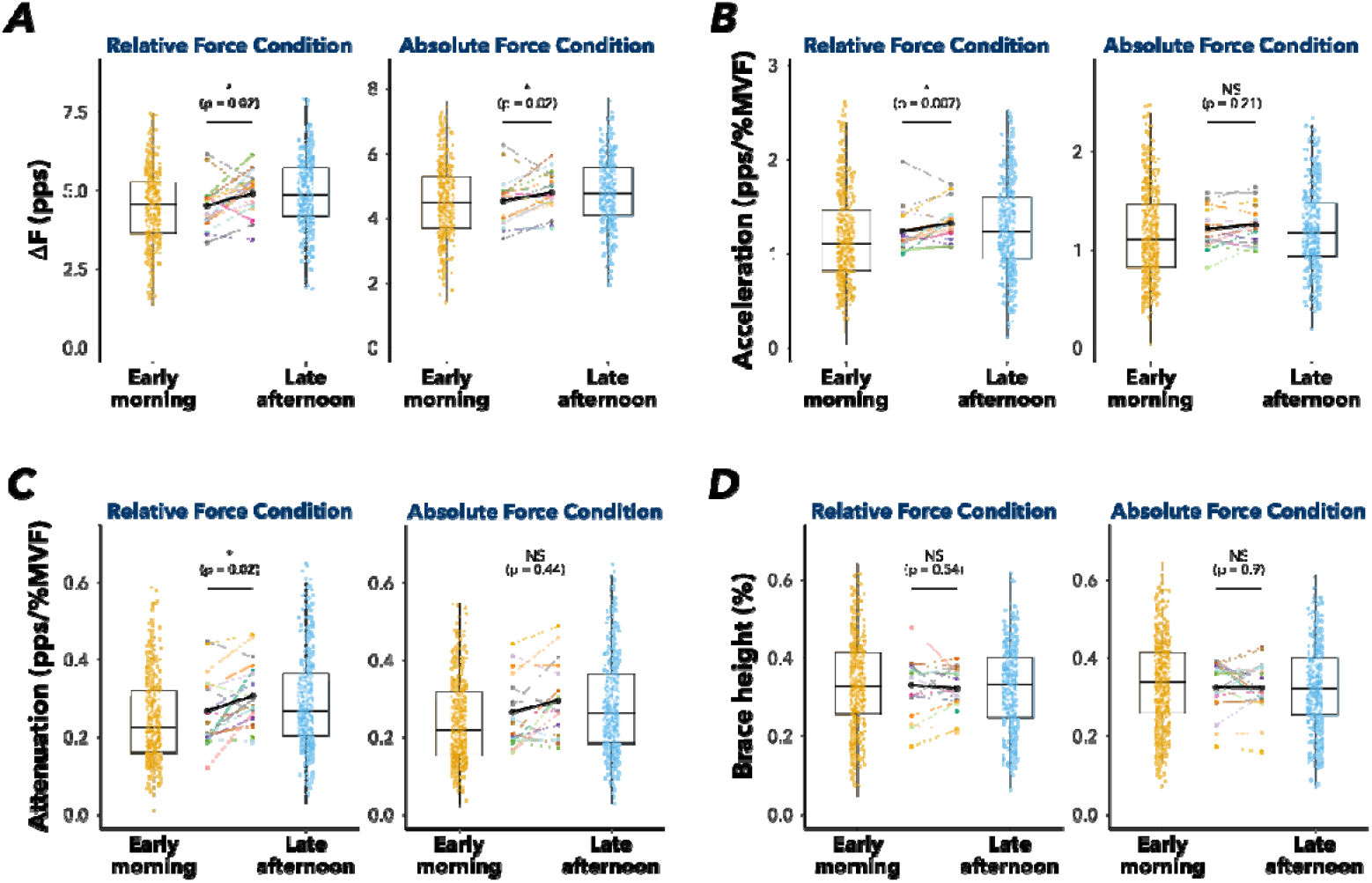
Changes in estimates of persistent inward currents from early morning to late afternoon (evening). ΔF estimates (**A**), acceleration (**B**), attenuation (**C**) and brace height (**D**) were obtained from the *tibialis anterior* (TA) muscle during early morning and late afternoon sessions. PIC prolongation effect was assessed using ΔF estimates. The geometric approach calculated brace height, acceleration, and attenuation, which were used to estimate the contribution of the neuromodulatory drive to motor unit discharge, persistent inward currents amplification effect on the discharge, and the pattern of inhibition during isometric dorsiflexion tasks, respectively. Tasks were conducted under two force conditions: (i) a relative force condition, set to 40% of the maximal voluntary force (MVF) measured during the same session (left panel), and (ii) an absolute force condition, set to 40% of the MVF recorded during the first visit (right panel), regardless of the time of day. Distributions of unit-wise ΔF values are illustrated using half-violin plots for each time-of-day condition. Boxplots represent the median (50^th^ percentile), lower (25^th^ percentile), and upper (75^th^ percentile) quartiles, with whiskers extending to 1.5 times the interquartile range. Colored dots represent individual participant means, and connecting lines indicate within-participant changes across conditions. Participant colours are consistent across all plots in the panel. Asterisks (*) indicate a significant main effect of time-of-day, with the alpha level set at 5%.

A significant time-of-day effect was also found in the absolute force condition (F(1, 17.27) = 7.255, p = 0.015), with ΔF values significantly 0.31 ± 0.23 pps greater in the late afternoon (4.8 ± 0.2 pps; 95% CI [4.4, 5.1]) compared to the early morning (4.4 ± 0.2 pps; 95% CI [4.1, 4.8]). This corresponded to a relative increase of +7.0% ± 5.1% (Figure 2).

In the relative force condition, changes in ΔF estimates between morning and late afternoon were moderately correlated with changes in MVF (r = 0.53, 95% CI [0.1, 0.8], p = 0.024) and with changes in peak discharge rate (r = 0.62, 95% CI [0.2, 0.8], p = 0.005). While both associations were supported by bootstrap-derived confidence intervals that excluded zero, the corresponding empirical p-values from bootstrap resampling were not statistically significant (p > 0.52 in both cases). This discrepancy suggests uncertainty in the robustness of these associations, likely driven by inter-individual variability and limited sample size. No significant associations were observed in the absolute force condition for either variable (p = 0.88 and 0.40, respectively).

### Quasi-geometric analyses of motor unit discharge trajectories

We also used a quasi-geometric method to assess PIC-related synaptic amplification and to better differentiate the respective contributions of neuromodulatory and inhibitory influences on PICs (Beauchamp et al., 2023). Focusing first on the acceleration phase, which is thought to reflect PIC amplification effect (Beauchamp et al., 2023), we observed a significant time-of-day effect in the relative (F(1, 18.21) = 9.144, p = 0.007) but not in the absolute force condition (F(1, 20.06) = 1.674, p = 0.21). In the relative force condition, acceleration values were significantly higher in the late afternoon (1.3 ± 0.04 pps/%MVF; 95% CI [1.2, 1.4]) compared to the early morning (1.2 ± 0.04 pps/%MVF; 95% CI [1.1, 1.3]), corresponding to a relative difference of +9.2% ± 4.6% (Figure 2).

The attenuation slope, which is presumed to reflect inhibitory inputs to motor neurons, exhibited a significant time-of-day effect in the relative (F(1, 19.24) = 6.722, p = 0.02), but not in the absolute force condition (F(1, 17.42) = 0.626, p = 0.44). In the relative force condition, attenuation values were 0.03 ± 0.03 pps/%MVF higher in the late afternoon (0.3 ± 0.02 pps/%MVF; 95% CI [0.3, 0.3]) compared to the early morning (0.26 ± 0.02 pps/%MVF; 95% CI [0.2, 0.3]). This corresponded to a relative increase of +13.4% ± 10.4% (Figure 2). Finally, brace height, a metric shown in simulation studies to be highly sensitive to neuromodulatory drive (Beauchamp et al., 2023; Chardon et al., 2024), showed no significant time-of-day effect in either the relative (F(1, 17.89) = 0.381, p = 0.54) or the absolute force condition (F(1, 18.06) = 0.016, p = 0.90).

No significant associations were observed between differences in discharge rate (both at peak and at recruitment) and differences in acceleration, attenuation, or brace height metrics between the two time points in either the relative or absolute force conditions (all p > 0.08).

## DISCUSSION

By analysing the discharge patterns of a large number of motor units (up to 92 per participant and condition) in the *tibialis anterior* muscle during submaximal contractions, we observed significantly greater PIC prolongation effect (*via* ΔF estimates) in the late afternoon compared to the early morning, regardless of the condition (relative or absolute contraction intensity). In addition, PIC amplification effect — estimated using the acceleration metric — was greater in the late afternoon during the relative force condition and was accompanied by a steeper attenuation slope. Taken together and considering the absence of difference in brace height between time points, these results suggest that the contribution of PICs to spinal motor neuron activity varies across the day and is likely driven by changes in inhibitory synaptic inputs rather than changes in neuromodulatory drive.

Neuromuscular performance, including maximal force production, is typically enhanced in the late afternoon compared to the early morning (Douglas et al., 2021; Knaier et al., 2022). While this diurnal pattern is well documented, the potential underlying neural mechanisms remain poorly understood. PICs, which amplify and prolong the effects of synaptic inputs to motor neurons, may play a key role in this process, yet their contribution has not been previously examined. Our findings indicate an enhanced contribution of PICs to motor unit behaviour in the late afternoon compared to the early morning. This was evidenced by (i) higher ΔF estimates in the late afternoon, regardless of force condition, indicating a greater prolongation effect of synaptic input, and (ii) a steeper acceleration of motor unit discharge rate in the relative force condition during the late afternoon, reflecting a stronger amplification effect. Although time-of-day differences in acceleration (approximately +9.2% under relative force conditions) and ΔF values (approximately +7.0-9.3% across conditions) were modest in magnitude, they are comparable to those observed in response to local vibration applied to the tibialis anterior (Lapole et al., 2023) or to changes in contraction intensity (Mackay et al., 2023). Overall, it suggests that these time-of-day effects may significantly influence motor unit behaviour. While it might be tempting to attribute the greater maximal torque observed in the late afternoon to enhanced PIC effects on motor unit activity, it is important to note that PIC-related metrics in our study were assessed during submaximal contractions. Thus, while our findings support time-of-day–dependent modulation of PIC contribution under submaximal conditions, they do not allow us to directly infer effects on maximal torque production. Of note, ramp contractions up to MVC could not be performed without inducing fatigue, which would have compromised the accuracy of PIC estimates. Despite these limitations, our findings provide strong evidence that motor neuron excitability (as inferred from PIC-related metrics) is enhanced during submaximal contractions in the late afternoon compared to early morning. Consistent with the findings of (Hirono et al., 2024), we observed higher discharge rates but unchanged recruitment thresholds in the relative force condition in the late afternoon, suggesting an altered capacity to modulate discharge rate across the day. This pattern likely reflects increased motor neuron excitability, enabling higher firing rates for a given synaptic input.

Given that PIC activation is modulated by both neuromodulatory and inhibitory inputs to motor neurons, we used complementary metrics proposed by Beauchamp et al. (2023) to estimate the relative contributions of these inputs to motor unit discharge behaviour. We found that brace height — a metric identified in simulation studies as a robust indicator of neuromodulatory drive (Beauchamp et al., 2023; Chardon et al., 2024) — did not differ between early morning and late afternoon sessions. This finding suggests that the observed increase in PIC contribution to motor neuron behaviour in the late afternoon is unlikely to be driven by augmented monoaminergic (neuromodulatory) input to spinal motor neurons. Although this may appear to contradict some earlier findings of higher serotonin levels in the late afternoon or evening on brain samples (Asano, 1971; Carlsson et al., 2007; Matheson et al., 2015), it is important to note that there is no evidence of diurnal variation in spinal serotonin level in humans (Anand et al., 2024). Another possible explanation for the observed changes in PICs involves diurnal variation in inhibitory drive to the motor neurons. Evidence from animal and computational studies has consistently demonstrated the inhibitory dampening effect on PICs (Bui et al., 2008; Hyngstrom et al., 2008; Kuo et al., 2003), a finding corroborated by recent studies in humans (Mesquita et al., 2022; Orssatto et al., 2022; Pearcey et al., 2022; Revill & Fuglevand, 2016). Although the inhibitory drive to spinal motor neurons was not directly quantified in our study, we estimated the inhibitory-excitatory influences on motor unit discharge behaviour *via* measures of the attenuation phase (Beauchamp et al., 2023; Chardon et al., 2024). We observed a significant time-of-day effect on attenuation in the relative force condition, suggesting a modulation of PIC contribution to motor neuron activity through reduced inhibitory synaptic inputs in the late afternoon. This finding partly aligns with previous study reporting increased efficacy of transmission between Ia afferent fibres and spinal motor neurons in the evening compared to the morning (Lang et al., 2011). However, these observations remain debated (Brangaccio et al., 2024), and other mechanisms such as pre-synaptic inhibition may also contribute to the observed effects. Together, our findings support the hypothesis that diurnal increases in PIC contribution to motor unit behaviour observed in the late afternoon relative to the early morning, are more likely mediated by reduced inhibitory input than by increased neuromodulatory drive to spinal motor neurons.

Several methodological aspects require consideration. First, we were unable to track a substantial number of motor units across the early morning and late afternoon sessions. Despite using validated tracking methods, such as the cross-correlation of motor unit action potentials (Martinez□Valdes et al., 2017) or the re-application of motor unit filters (Francic & Holobar, 2021), the success rate remained low, with an average of 4.6□±□7.9 matched motor unit pairs per participant (range: 0–31, total of 82 motor units considering both force conditions) using the cross-correlation (threshold ≥ 0.8) of motor unit action potentials. This limited tracking efficiency is likely due to minor differences in electrode repositioning between sessions (despite careful electrodes marking procedures on the skin) and, more critically, to the non-stationary nature of the force profile (triangular ramps), which may alter motor unit filter characteristics over time. However, given the rigid control of motor units governed by the size principle (Henneman, 1957), it is unlikely that entirely different motor unit with different characteristics were active across sessions. Importantly, by decomposing EMG signals from 256 electrodes, we were able to identify a large number of motor units per participant (up to 92). As a result, our findings are based on representative motor unit populations, which gives us confidence that the inability to track individual motor units across sessions does not affect the validity of our conclusions. Second, we examined time-of-day difference in estimates of PICs on only two separate occasions: once in the morning (between 7:00 and 8:30 a.m.) and once in the late afternoon on a different day (between 5 and 7:30 p.m.). These time points were selected as they represent the bathyphase and acrophase of neuromuscular performance, respectively (Davison, 2022). While this study design enables meaningful comparison across distinct periods, it does not capture the full diurnal profile of PIC contribution to spinal motor neuron activity. Given that neuromodulatory and inhibitory inputs may vary more gradually or non-linearly throughout the day, additional time points would be needed to fully characterize the diurnal dynamics of PICs. Finally, interindividual variability in PIC contribution to motor unit behaviour and its diurnal modulation may also contribute to the observed patterns. Larger, more targeted studies are needed to address this question.

In conclusion, this study provides evidence for time-of-day–dependent variations in the contribution of PICs to motor unit discharge behaviour, highlighting that intrinsic motor neuron excitability (and thus spinal gain modulation) fluctuates across the day. These findings contribute to a deeper understanding of the temporal (diurnal) dynamics underlying the neural control of muscle contraction. They also underscore the importance of considering time-of-day as a potential confounding factor in experimental protocols investigating motor unit behaviour and PIC-related mechanisms. Further research is needed to clarify the underlying mechanisms and to determine how these diurnal fluctuations interact with task demands, fatigue, and pathological conditions.

## ACKNOWLEDGEMENTS

The authors would like to thank Alyssa Mannent for her valuable assistance during part of the data collection sessions, and François Dernoncourt for his support in data analysis.

## Notes

**Data availability:** The data that support the findings of this study are available on request from the corresponding author.

**Conflict of interests:** The authors declare no competing financial interests.

**Funding resources:** This work was supported by Université Côte d’Azur (CSI program) and by the French government, through the UCAJEDI Investments in the Future project managed by the National Research Agency (ANR) with the reference number ANR-15-IDEX-01, and by an ANR grant (ANR-24-CE17-5805, NEUROMOTOR project).

### Competing Interest Statement

The authors have declared no competing interest.

